# KAT3 mutations impair neural crest migration through EMT regulators *snai1b a*nd *snai2* in Rubinstein Taybi Syndrome

**DOI:** 10.1101/2024.05.19.593474

**Authors:** Shweta Verma, Sujit Dalabehera, Subhash Gowda, Koushika Chandrasekaran, Dayanidhi Singh, Bhavana Prasher, Sharmila Bapat, Sivaprakash Ramalingam, Chetana Sachidanandan

## Abstract

Rubinstein Taybi syndrome, a rare congenital disease is caused by mutation in KAT3 genes, *EP300* and *CREBBP*. A subset of tissues affected in RSTS have their origin in neural crest cells, prompting our exploration into the role of KAT3 in neural crest development. Our zebrafish RSTS models generated by knocking down or mutating *ep300a* and *cbpa* genes, reveal defects in neural crest migration and its derived tissues when KAT3 genes are perturbed. We also demonstrate that the effects on neural crest can be reversed by HDAC inhibition in in morphant embryos. KAT3 knockdown causes downregulation of EMT regulators, *snai1b* and *snai2*. Snai2 is known to repress *cdh6b* in neural crest cells facilitating their delamination from neural tube and migration. We generated RSTS patient-derived iPSC line and differentiated them into neural crest cells in vitro. We show that role of KAT3 proteins in neural crest migration is conserved in human iPSC derived neural crest cells. Our findings make a case for classifying RSTS as a neurocristopathy.

**Highlights:**

- Perturbation of KAT3 gene expression in zebrafish recapitulates the Rubinstein Taybi patient defects
- The zebrafish model of Rubinstein Taybi model reveals defects in neural crest cell migration
- KAT3 proteins regulate *snai2, snai1b* and cdh6, genes important for neural crest migration
- The neural crest migration defects in the zebrafish model can be partially rescued by modulating the global acetylation levels
- Study of RSTS patient-derived neural crest cells reveals that the role of KAT3 in neural crest migration is conserved across vertebrates

**Graphical Abstract:** 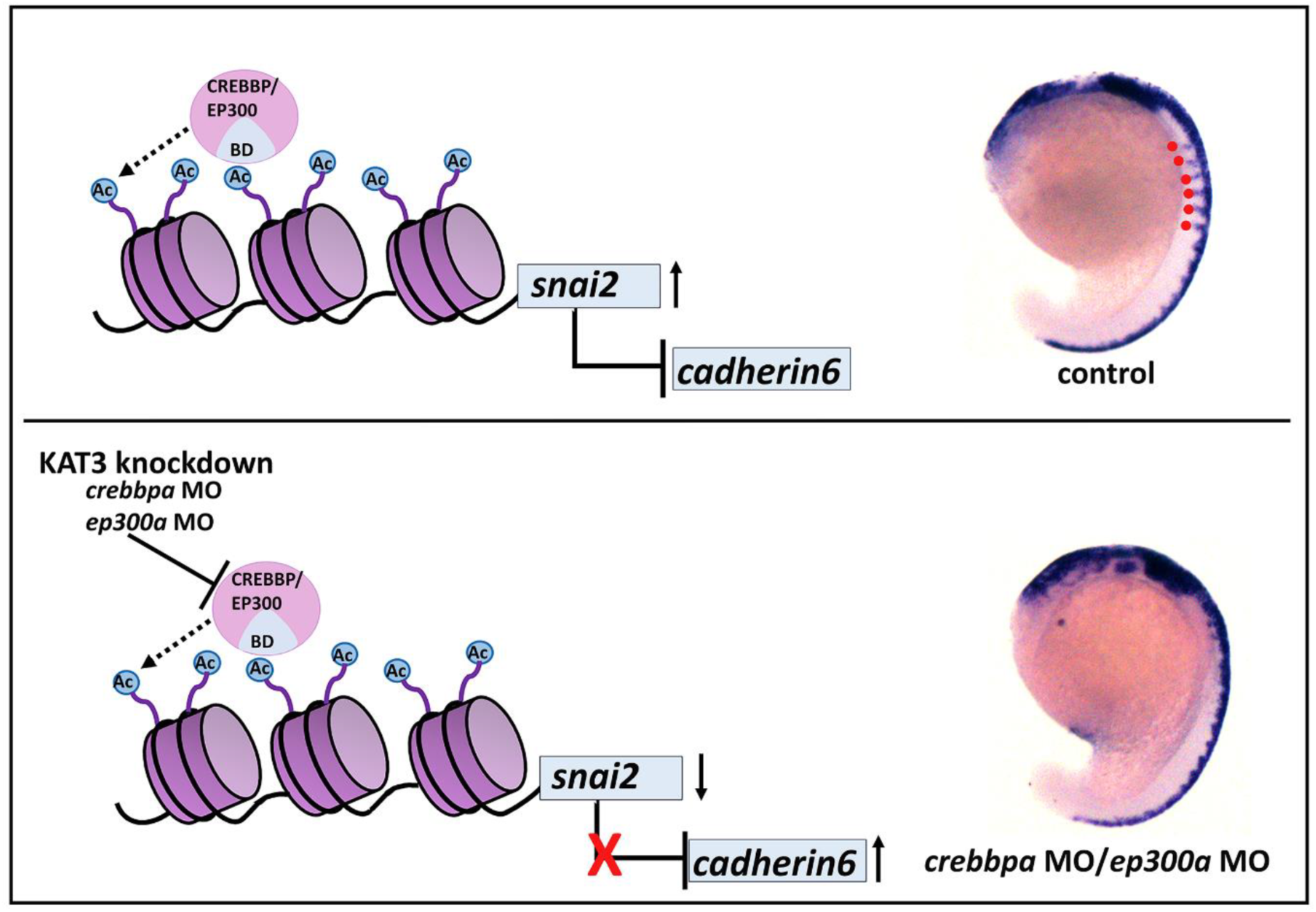

## 1. Background

Rubinstein Taybi Syndrome (RSTS) is a rare autosomal dominant congenital disorder (RSTS; OMIM #180849, #613684) caused by mutations in lysine acetyltransferase type 3 (KAT3) proteins, CREBBP and EP300 (1–3). The syndrome is characterized by broad thumb and halluces, growth retardation, heart defects and severe to moderate intellectual disability (4). The patients have distinct craniofacial features (5,6) and occasionally suffer from severe constipation caused by defects in the enteric nervous system (7–10). A number of these defects in RSTS can be traced to tissues that originate from neural crest cells suggesting that patients may have defects in neural crest cells during development. However, no role for *EP300* or *CREBBP* have been reported in neural crest development.

We have previously developed a zebrafish model for RSTS. The zebrafish embryos and larvae with *ep300a* knocked down showed craniofacial, limb and heart defects reminiscent of defects seen in patients (11). We have demonstrated that these defects can be rescued by elevating global histone acetylation levels either by inhibiting HDACs or by overexpressing the histone acetyl transferase (HAT) domain of Ep300a in zebrafish embryos(11).

Here, we illustrate that chemical inhibition of KAT3 proteins as well as knockdown of *crebbpa* and *ep300a* in zebrafish lead to defects in neural crest migration and neural crest derived tissues such as craniofacial cartilage, heart and enteric neurons. Similar migration defects are also observed in neural crest cells derived from a RSTS patient iPSCs (IGIBi018-A). We show that putative EP300 target genes *snai1b* and *snai2* are downregulated in the KAT3 knocked down zebrafish embryos, which in turn leads to upregulation of *cdh6* expression, likely preventing delamination and migration of neural crest cells.

## 2. Results

### 2.1 KAT3 regulates neural crest development in zebrafish

Patients with Rubinstein Taybi Syndrome have defects in tissues derived from neural crest such as craniofacial dysmorphism, gastrointestinal defects and cardiovascular defects (4,5,7–9). Therefore, we hypothesized that the KAT3 histone acetyl transferases would have a role in neural crest development. To test this, we inhibited the zebrafish KAT3 proteins Crebbp and Ep300 using the competitive inhibitor C646 (12). Embryos were exposed to 5µM C646 from 8 hpf (hours post fertilization) and the expression of *sox10*, the pan neural crest marker was analysed by RNA in situ hybridization at various stages of development. At 3ss (somite stage), when the neural crest cells (NCCs) first appear, *sox10* expression was evident at the neural plate border in control embryos whereas in the C646 treated embryos, it was less apparent and further apart (Fig.1A-B). At 6ss, NCCs begin medial migration, which was absent in C646 treated embryos (Fig.1C-D). NCCs first emerge in the rostral region followed by more caudal regions. By 10ss, NCCs were evident in the rostral three-quarters of the embryos in control whereas when KAT3 was inhibited *sox10* expression became restricted to the rostral half of the embryos (Fig. 1E-F). After NCCs emerge from the neural plate border they begin a dorsolateral migration forming streaks of migrating cells between the somites. At 18ss, this migration was prominent in the control embryos while was severely reduced in C646 treated (Fig. 1G-H). Thus, we observe a delay in neural crest development on chemical inhibition of KAT3 proteins.

**Figure 1:**
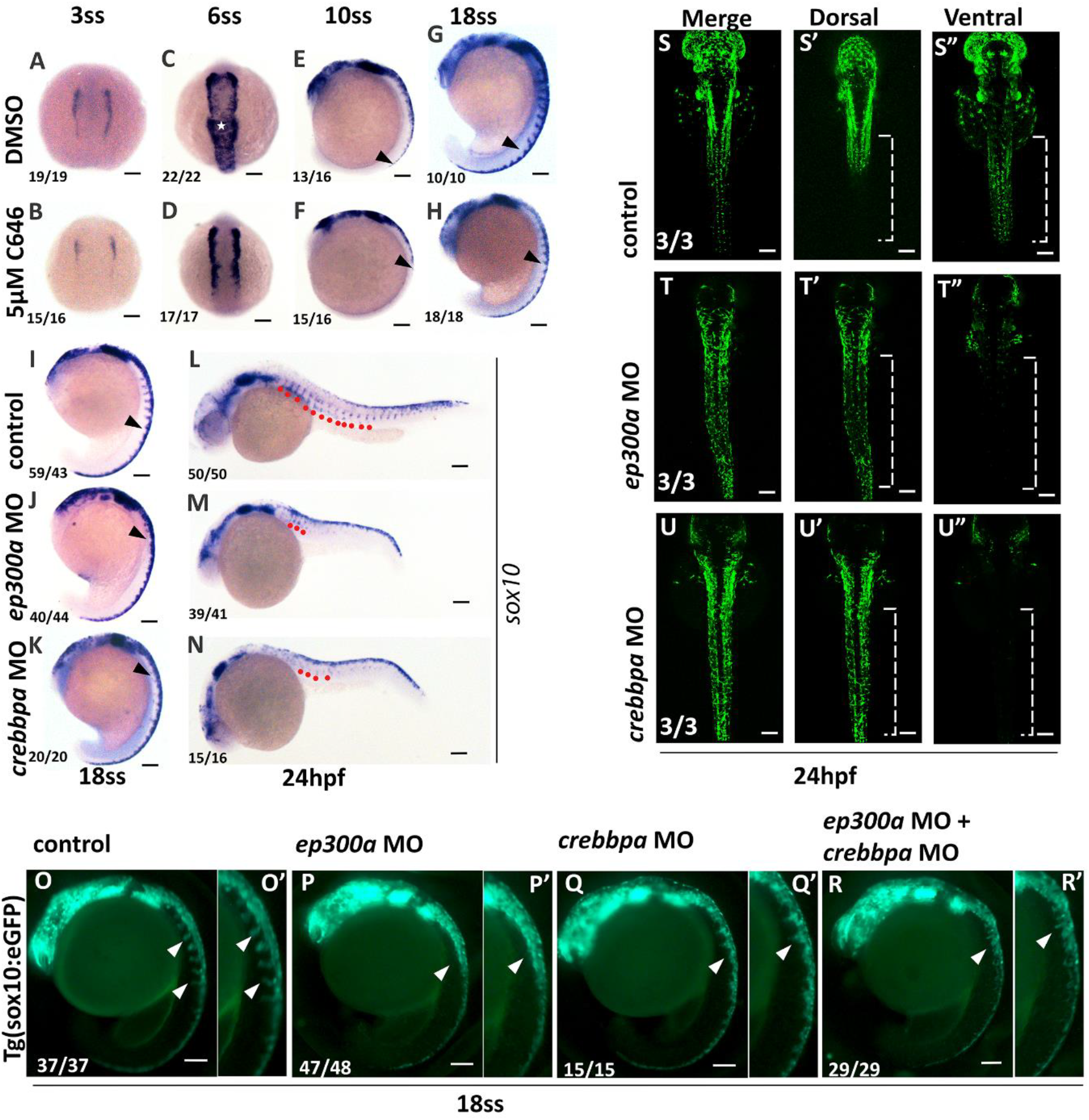
KAT3 regulates neural crest development in zebrafish. **(A-H)** RNA in situ hybridization of *sox10* at different developmental stages of DMSO control (A, C, E, G) and C646 treated zebrafish embryos (B, D, F, H). (A, B) At 3 somite stage(ss), expression of *sox10* is apparent at the neural plate border. In C646 treated embryos the sox10 expression marks only shorter strip of neural plate border. (C, D) At 6ss, neural crest shows medial migration (white asterisk) in control whereas the crest does not appear to be migrating medially in C646 treated embryos. (E, F) At 10ss, NCCs in C646 treated embryos have emerged in half of the length of the embryo (black arrowhead) while in control NCCs are evident in about two-thirds of the length of the embryo. (G, H) At 18ss, dorsoventral migration of NCCs (black arrowhead) marked by sox10 expression is less evident in C646 treated embryos compared to controls. **(I-N)** RNA in situ hybridization of *sox10* at 18ss and 24hpf in *ep300a* and *crebbpa* morphants. (I, J, K) At 18ss, dorsoventrally migrating NCCs are visible as streams between somites (black arrowhead) in control embryos whereas there is much less migration in *ep300a* and *crebbpa* morphants. (L, M, N) At 24hpf, NCCs have migrated towards the ventral side of the embryo (red dots) whereas in *ep300a* and *crebbpa* morphants NCCs are stalled near the dorsal side of the embryos (red dots). **(O-R)** At 18ss, *Tg (sox10* : *eGFP)* marks the NCCs which show dorsoventral migration (white arrowheads) in the controls (O, O’) while the migrating streams of cells are absent in *ep300a* (P,P’), *crebbpa*(Q, Q’) and double morphants (R, R’). (O’, P, Q, R’) are magnified images of (O, P, Q, R). **(S-U”)** Confocal projections of neural crest cells in different planes of *Tg (sox10* : *eGFP)* embryos. Sox10 expressing trunk NCCs are predominantly seen in the ventral projections as the cells have migrated ventrally in control embryos (S, S’, S”). *ep300* (R, T’, T”) and *crebbp* (U, U’, U”) morphants show NCCs more in the dorsal projection than the ventral because of impaired dorso-ventral migration. (A-D)(S-U”) Dorsal views, anterior to the top. (E-H)(I-K)(O-R’) lateral views, anterior to the top, dorsal to the right. (L-N) lateral views, anterior to the right, dorsal to the right. Scale bar = 100µm. Numbers at the bottom left corner represent number of embryos showing particular phenotype over the total number of embryos imaged.

Rubinstein Taybi Syndrome is caused by mutations in either *CREBBP* (KAT3A) or *EP300* (KAT3B) genes. We investigated the individual roles of KAT3A and KAT3B in neural crest by knocking these genes down in zebrafish. Zebrafish has two co-orthologs of *EP300* and *CREBBP* genes each namely, *ep300a* and *ep300b*, and *crebbpa* and *crebbpb* respectively (11). Our previous studies showed that knockdown of *ep300a* recapitulates RSTS patient phenotypes in zebrafish embryos. To model RSTS type 1 (KAT3A mutation) in zebrafish, we knocked down the expression of *crebbpa* and *crebbpb* using splice block antisense morpholino oligonucleotides (MO). One-cell stage embryos were injected with 4ng of *crebbpa* MO or 8ng of *crebbpb* MO (detailed methodology in supplementary Fig. S1). We observed developmental defects including jaw defects, heart edema and smaller eyes in *crebbpa* knocked down embryos at 4 dpf (Supplementary Fig. S1). *crebbpb* MO injected embryos displayed no overt developmental defects. Both MOs caused splicing defects as confirmed by PCR (Supplementary Fig.S1). As reported previously, we have also found that knockdown of *ep300a* but not *ep300b* by splice-block antisense MO led to developmental defects in zebrafish embryos at 4dpf (11) (Supplementary Fig. S1). Therefore, all further experiments in this study were performed using only *ep300a* and *crebbpa* MOs.

To address the role of *ep300a* and *crebbpa* in neural crest, we performed RNA in situ hybridization for *sox10* at different developmental stages. Expression pattern of *sox10* in *ep300a* and *crebbpa* knocked down embryos at 3ss, 6ss and 10ss were unaffected compared to control embryos (Fig. S2 A-I). However, the dorsolateral migration of neural crest cells evident at 18ss was significantly diminished in embryos injected with MOs against *ep300a* and *crebbpa* (Fig. 1I-K). Similar defects were also observed using *crestin*, another marker of neural crest cells (Fig. S2 J-M). Expression of *sox10* transcription factor is downregulated in the NC cells as they complete their dorsolateral migration. In *ep300a* and *crebbpa* morphant embryos the persistence of the *sox10* signal in the dorsal NC cells of the 24 hpf embryos indicates defect in the migration of NC cells (Fig. 1L-N). We also visualized this migration in *Tg(sox10* : *eGFP)* transgenic line with fluorescently labelled neural crest cells. At 18ss, migrating neural crest cells were absent in the *ep300a* and *crebbpa* morphants (Fig. 1O-R). In 24 hpf *ep300a* and *crebbpa* morphant embryos neural crest cells remain on the dorsal side of the embryo and we observe very little dorsolateral migration (Fig. 1S-U”).

### 2.2 Neural crest derived tissues are affected in KAT3 morphants

Neural crest cells are multipotent cells that differentiate into multiple cell types including melanocytes, chondrocytes, glia, neurons, osteocytes and cardiomyocytes. Since the correct migration of neural crest cells is essential for their differentiation into various tissue types, we analysed the tissues derived from neural crest cells in *ep300a* and *crebbpa* morphants. Embryos injected with either *ep300a* or *crebbpa* MO have reduced lower jaw (Fig. 2A-C). We observed MO injected *Tg(sox10:eGFP)* embryos at 4dpf and found an absence of Meckel’s cartilage and a lack of demarcation of the ceratobranchial cartilages (Fig. 2D-F). Ablation of neural crest has been reported to cause defects in the development of the heart structure and function in zebrafish (13). The *ep300a* and *crebbpa* morphants show severe pericardial edema suggesting heart defects (Fig. 2A-C). We injected the MOs in *Tg(myl7: RFP)* transgenic zebrafish line to observe the heart structure. Ventricular size and the heart tube folding were both defective in the *ep300a* and *crebbpa* morphants compared to the control (Fig. 2J-L).

**Figure 2:**
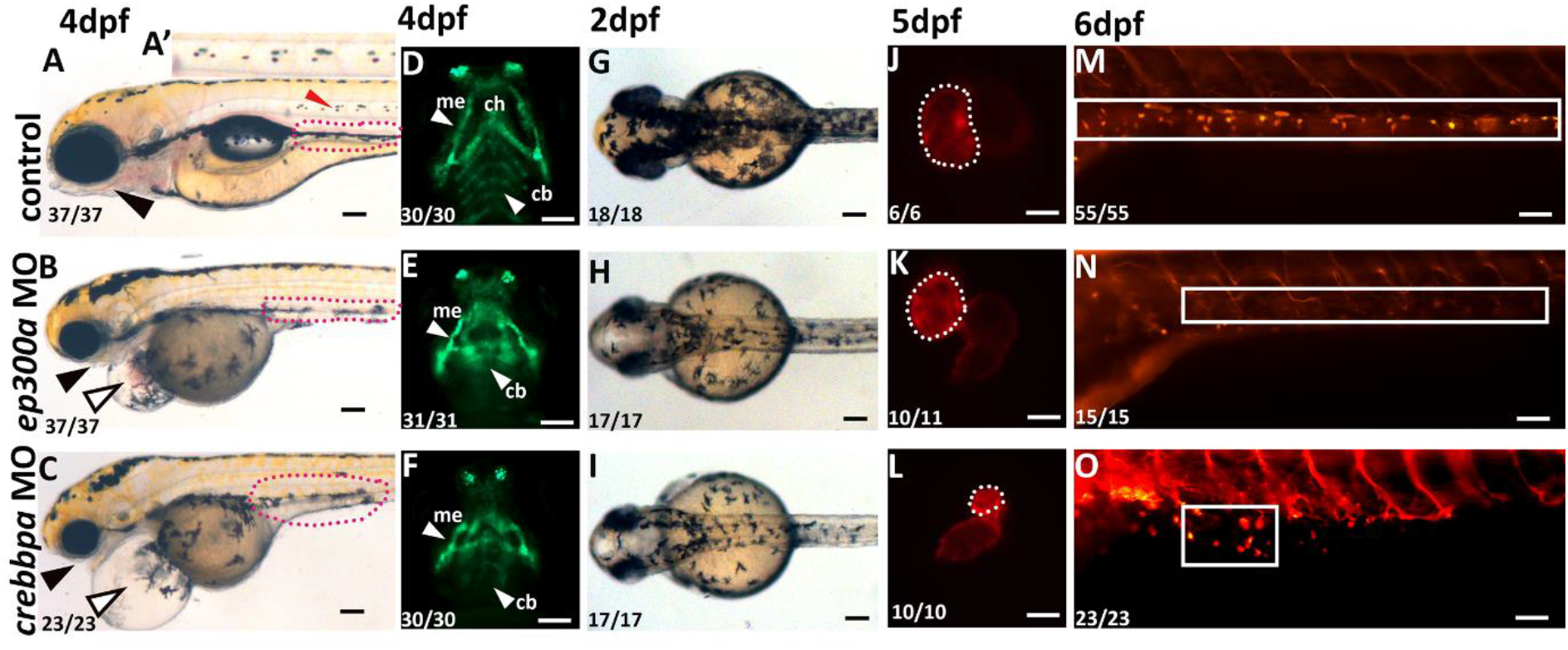
Neural crest derived tissues are affected in KAT3 morphants. **(A-C)** At 4dpf, lower jaw is well defined in control embryos (black filled arrowhead) while in the *ep300a* and *crebbpa* morphants it is absent. (A) Melanophores form ventral stripe (red dots) and lateral stripe (red arrow head). A’ is magnified image of the lateral stripe in control embryo. (B, C) *ep300a* and *crebbpa* morphants lack lateral and ventral melanophore stripes. These morphants have heart edema as well (black empty arrowhead). **(D-F)** At 4dpf, *Tg(sox10* : *eGFP)* marks craniofacial cartilages such as Meckel’s cartilage (me), ceratohyal cartilage (ch) and ceratobranchial cartilages (cb). *ep300a* and *crebbpa* morphants have shorter Meckel’s and Ceratohyal cartilage, and ceratobranchial cartilages are diffused (white arrowhead). **(G-I)** At 2dpf, control embryos have higher melanophores compared to *ep300a* and *crebbpa* morphants. **(J-L)** At 5dpf, *Tg(myl7:RFP)* is used to visualize the heart. Compared to the control heart with a well-developed heart with trabeculated ventricle (marked by dotted white line) the *ep300a* and *crebbpa* morphant hearts show looping defects, reduced ventricles with irregular trabeculation. **(M-O)** At 6dpf, enteric neurons visualized by *Tg(NBT:dsRed)* were spread along the entire length of the gut in the control embryos. *ep300a* morphants show an absence of enteric neurons whereas in *crebbpa* morphants there were fewer enteric neurons and these were unorganized (white box). (A-C)(M-O) lateral views with anterior to the left. (D-F) (J-L) Ventral views, anterior to the top. (G-I) Dorsal views with anterior to the left. Scale bar = 100µm. Numbers at the bottom left corner represent number of embryos showing particular phenotype over the total number of embryos imaged.

Melanophores are one of the cell types originating from neural crest and *ep300a* and *crebbpa* MOs have noticeable impact on melanophores. Melanophores are decreased in number in 2dpf *ep300a* and *crebbpa* morphants (Fig. 2G-I) and show defective migration in 4dpf (Fig. 2A-C).

The role of KAT3 proteins has not been studied in enteric neuron development. However, severe and chronic constipation, a hallmark of enteric neuron defects, is a common feature of Rubinstein Taybi Syndrome (5,14). Enteric neurons are derived from migratory vagal neural crest cells and we tested the effect of KAT3 knockdown in *Tg(NBT:dsRed*) transgenic zebrafish line. The enteric neuron cell bodies were well articulated in 6dpf control larvae, but in the *ep300a* and *crebbpa* morphants these cells were either absent or aberrantly placed respectively (Fig. 2M-O).

### 2.3 EP300 regulates the expression of genes essential for neural crest

To identify KAT3 dependent neural crest regulators we used publicly available data. Wysocka and group have performed an EP300 Chip-Seq to identify enhancers active in human stem cell derived Neural Crest Cells (hNCC) with human Embyonic Stem cells (hESC) as controls (15). We used this data to identify potential target genes of EP300 in the neural crest. Enhancer coordinates bound by EP300 were assigned to genes within 100kb of TSS using the PAVIS software (Supplementary Data 1) (16). This approach identified 2462 potential targets bound by EP300 in neural crest cells. To identify genes expressed in neural crest, we analysed the RNA-Seq data in the same hNCC compared to human neuro-ectodermal cells (hNEC)(15). We shortlisted 480 genes that showed 2-fold increase in expression in hNCC over hNEC (Supplementary Data 2). The two lists generated were compared and 121 common genes were identified (Fig. 3A). From these 121, we shortlisted 10 genes that have known roles in neural crest development (Supplementary Data 3).

**Figure 3:**
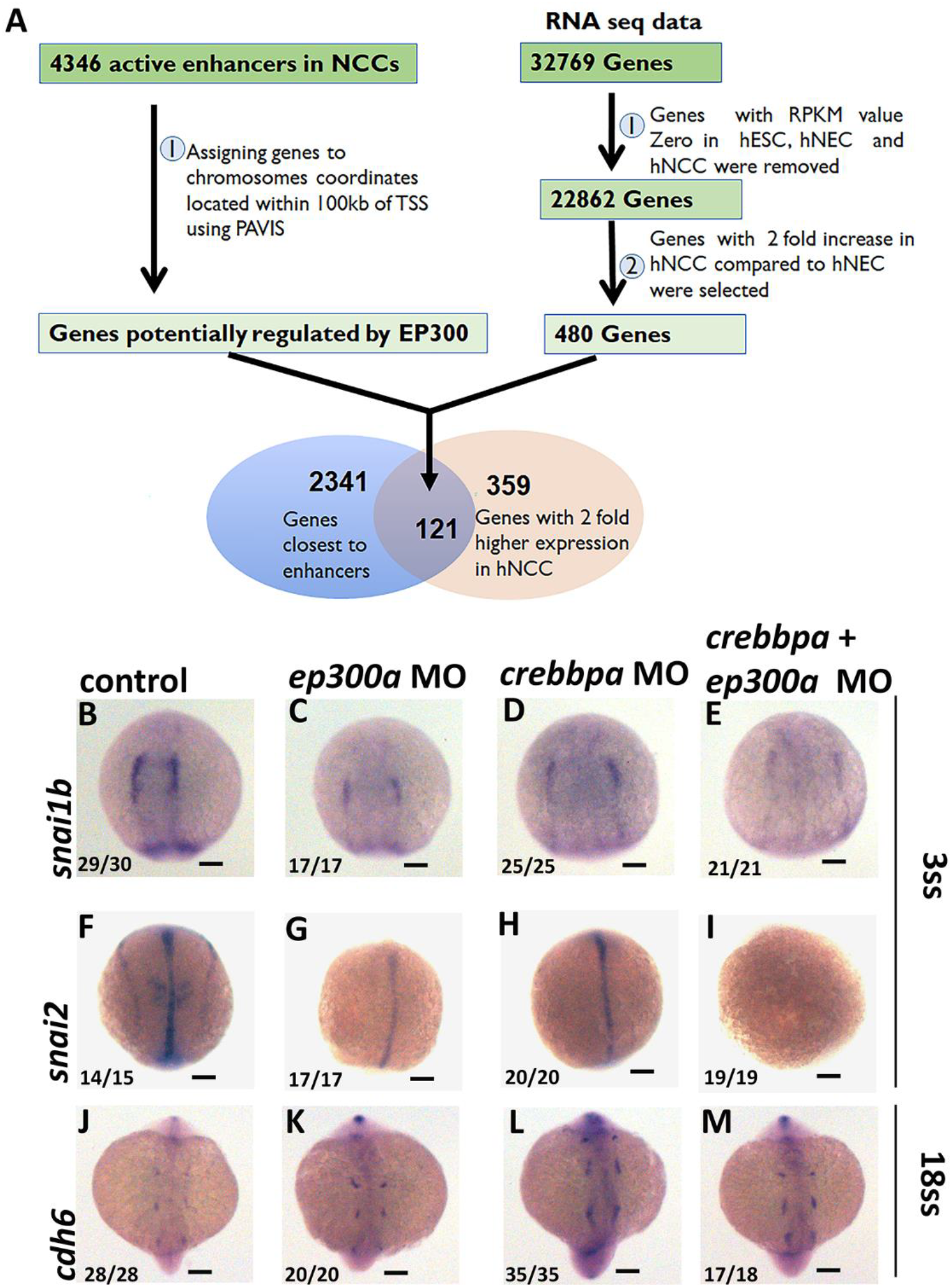
EP300 regulates the expression of genes essential for neural crest. **(A)** Schematic for the workflow to identify EP300 target genes with NC function. 4346 enhancers were occupied by EP300 as identified in the publicly available ChIP seq data from human stem cell derived NCC. 2341 genes were assigned to active enhancer chromosomal coordinates (bound by EP300) within 100kb of TSS using PAVIS online tool. From the 32769 genes identified in the publicly available RNA seq data of human ESC derived NCC 480 genes showed 2-fold upregulation in NCC. 121 putative Ep300 target genes were identified by overlapping the RNA seq and ChIPseq genes. **(B-I)** RNA in-situ hybridisation of putative target genes reveals reduced expression of *snai1b* and *snai2* in *ep300a* and *crebbpa* morphants at 3ss. **(J-N)** RNA in-situ hybridisation indicates upregulation of expression of *cadherin6* (*cdh6*) at 18ss. Dorsal views with anterior to the top. Scale bar = 100 µm. Numbers at the bottom left corner represent number of embryos showing particular phenotype over the total number of embryos imaged.

We performed RNA in-situ hybridization against these genes in control and KAT3 knockdown embryos. *dlx2a, pax3, msxb, tfap2a, sox8*, and *mycn* showed no difference in expression between control and *ep300* or *crebbp* morphants (Supplementary Fig. S3 A-F’). At 18ss, expression of *lef1* and *meis2a* was absent or downregulated in *ep300*a morphants in comparison to the control embryos (Supplementary Fig. S3 G-H’). *lef1* and *meis2* are required for the development of neural crest derived cranial and cardiac neural crest cell formation (17) as well as for tooth morphogenesis (18). Although lef1 and meis2 have been shown to be important for melanocyte differentiation and cranial neural crest development respectively (19,20), there is currently no report of their having a role in neural crest migration. This needs further investigation. We observed downregulation of the crest expression of *snai1b* and *snai2* in 3ss embryos injected with *ep300* and *crebbp* morpholinos (Fig. 3B-I). Downregulation of *cdh6* is known to be essential for epithelial to mesenchymal transition of neuroectodermal cells to form neural crest (21). We observed an upregulation of *cdh6* in KAT3 morphants compared to control embryos suggesting a role of KAT3 proteins in regulating neural crest delamination (Fig. 3J-M).

### 2.4 *ep300a* ^*+/*BD1Δ^ mutants have defect in migration of neural crest cells

Efforts to knockout *EP300* have been made in various organisms and have failed due to embryonic lethality (22,23). We tried to create an *ep300* mutant using CRISPR/Cas9 technology in zebrafish. We used a guide RNA designed against the bromodomain of Ep300a and identified putative germline mutations in an F0 fish after numerous efforts. The sequencing of the target region in the F2 generation revealed a heterozygous allele with a single-base deletion, predicted to result in a truncated Ep300a protein (Fig. 4A-C). This mutant allele will be called *ep300a* ^*+/*BD1Δ^ henceforth. We observed severe developmental defects in the mutants including smaller head size, jaw defect, smaller eyes and heart edema at 4dpf (Fig. 4D-E). We observed a decrease in the melanophore pigmentation in *ep300a* ^*+/*BD1Δ^ mutants in comparison to the control embryos at 2dpf (Fig. 4F-G). We crossed the F1 *ep300a* ^*+/*BD1**Δ**^ fishes to *Tg(Sox10:eGFP)* and at 4dpf, we observed that meckel’s and ceratohyal cartilages were shorter and wider in *ep300a* ^+/BD1Δ^ whereas the ceratobranchial cartilages were diffused in mutants compared to control (Fig. 4H-I).

**Figure 4:**
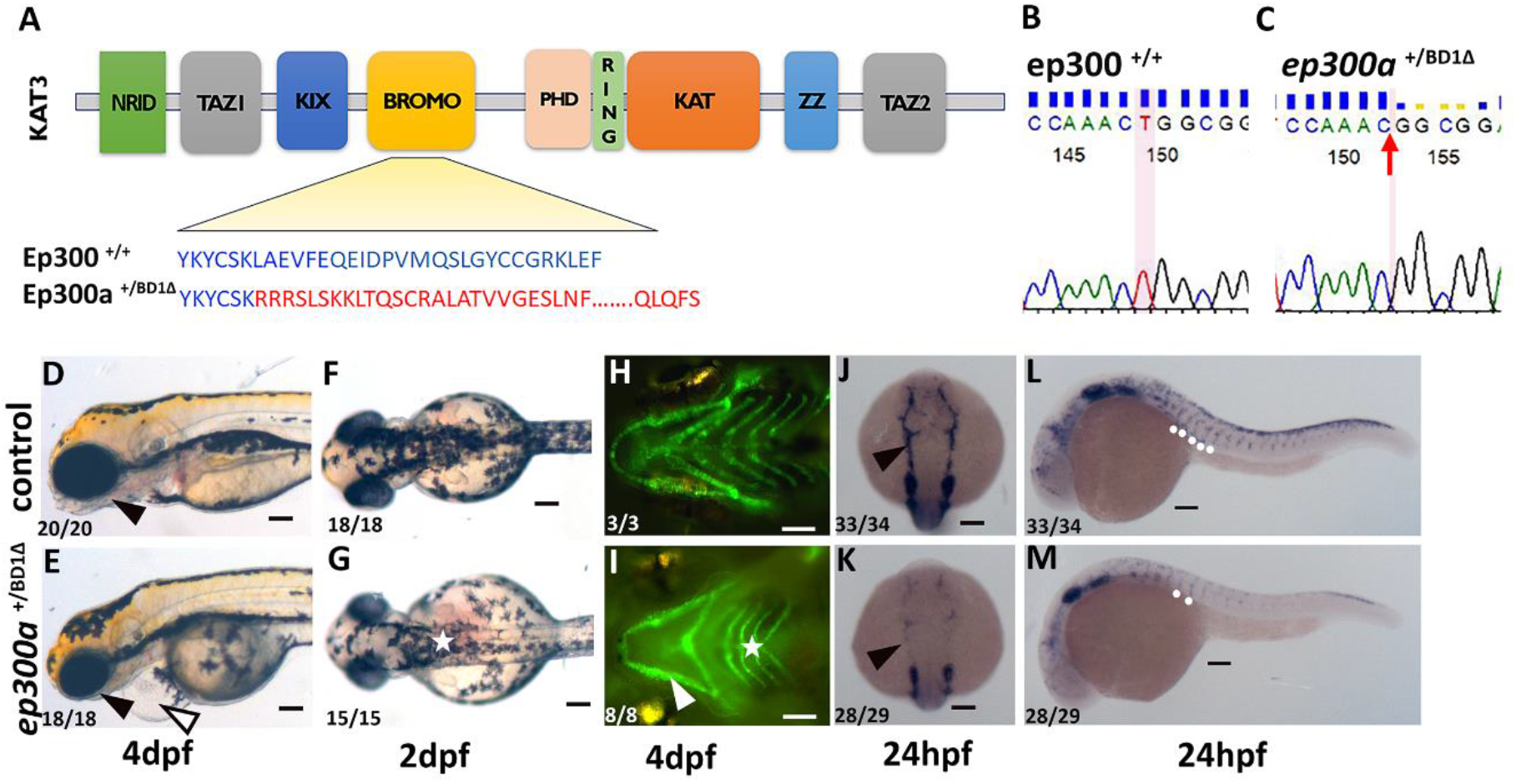
*ep300a* ^*+/*BD1D^ mutants have defect in migration of neural crest cells. **(A)** Schematic of KAT3 protein domains marking the bromodomain with mutation and the amino acid sequence of Ep300 and CRIPSR mutant *ep300a* ^*+/*BD1Δ^. **(B-C**) Chromatogram of ep300^+/+^ and *ep300a* ^*+/*BD1Δ^ indicating 1bp deletion (red arrow). **(D-E)** Brightfield image of 4dpf embryos from incross of *ep300a* ^*+/*BD1Δ^ show smaller eyes, heart edema, melanocyte defect and redundant jaw compared to control. **(F-G)** 2 dpf embryos from incross of *ep300a* ^*+/*BD1Δ^ display decrease in melanophores compared to control. **(H-I)** *Tg(sox10: eGFP)* crossed to *ep300a* ^*+/*BD1Δ^ displayed reduced craniofacial cartilages compared to control. **(J-M)** RNA in-situ hybridization reveals decrease in expression of *sox10* in cranial and trunk neural crest. (D-E)(L-M) Lateral views with anterior to the left. (F-G) Dorsal views with anterior to the left. (H-I) Ventral view with anterior to left. (J-K) dorsal view with anterior to top. Scale bar = 100 µm. Numbers at the bottom left corner represent number of embryos showing particular phenotype over the total number of embryos imaged.

Further, we looked at the expression of *sox10*, the neural crest marker, at 24 hpf in the heterozygous mutant embryos. The cranial NCCs that are derived from the migration of rostral NCCs showed a significant reduction (Fig. 4J-K). The trunk NCCs that emerge from the dorsal edge of the neural tube also showed significant reduction. The control embryos showed streams of neural crest cells migrating dorsolaterally between the somites and these were absent in the mutant embryos (Fig. 4L-M).

### 2.5 Modulating histone acetylation partially rescues neural crest defects

We have previously shown that inhibition of certain histone deacetylases partially rescues craniofacial cartilage defects in ep300 morphants (11). We performed a similar curated screen with HDAC and Sirtuin inhibitors to identify molecules that can rescue the neural crest migration defect observed in *ep300a* and *crebbpa* morpholino injected embryos and found three compounds with potential activity. DMSO treated *ep300a* and *crebbpa* morphants showed strong NCC migration defects (Fig. 5A, E). Morphant embryos treated with 10µM of CHIC35 and 5 µM of HDACi II, HDACiIII from 8hpf till 18hpf were analysed for *sox10* expression by RNA in-situ hybridization. We found partial rescue of dorsolateral migration of NCCs in *ep300a* morphants upon treatment with HDACi II, HDACi III and CHIC35 (Fig. 5B-D). NCC migration in *crebbpa* morphants could not be rescued by any of the compounds tested (Fig. 5F-H). This suggests a differential requirement for *crebbpa* and *ep300a* neural crest development.

**Figure 5:**
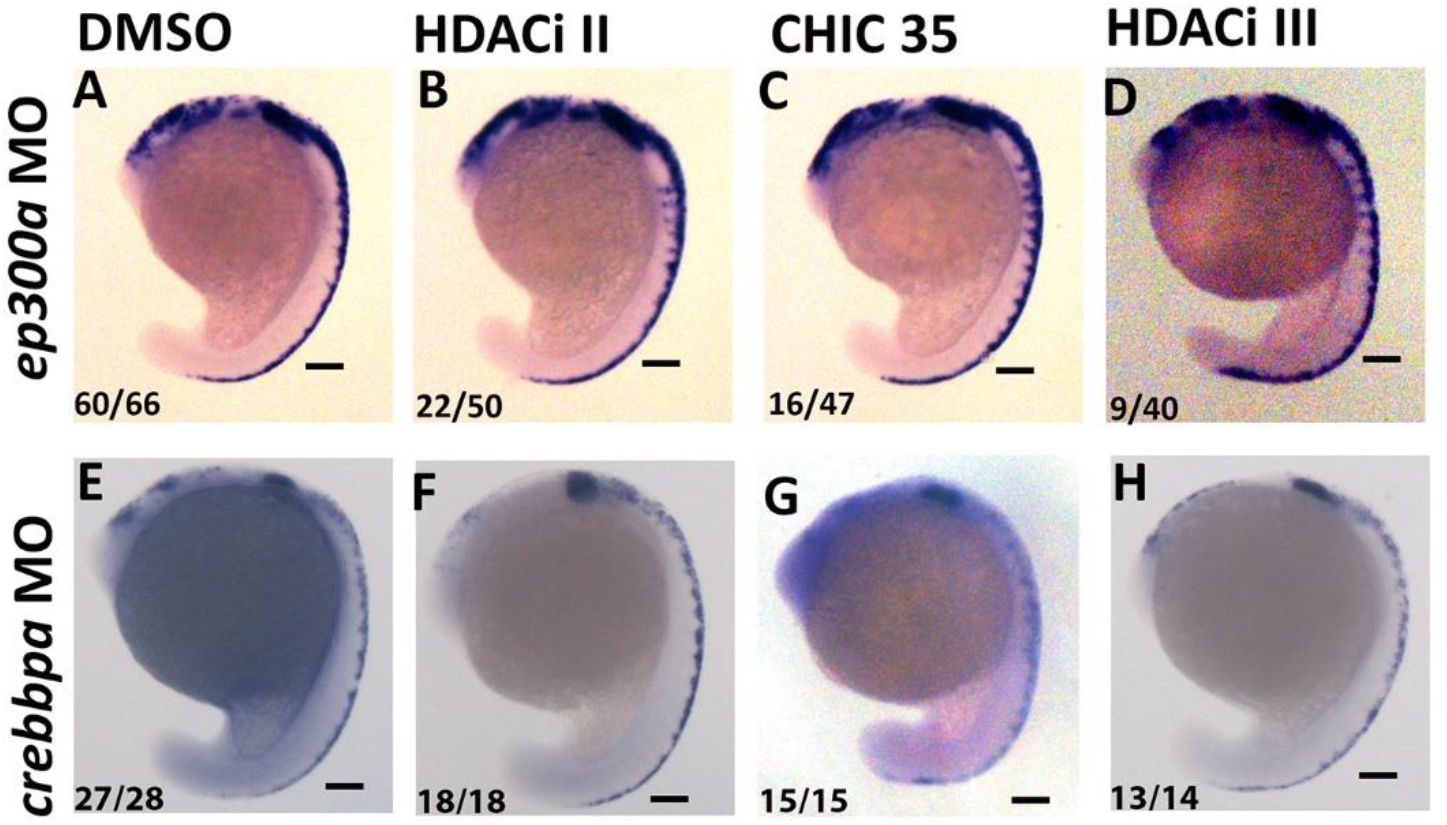
Modulating histone acetylation partially rescues neural crest defects. **(A-H)** RNA in-situ hybridization of *sox10* shows partial rescue of dorsolateral migration of neural crest cells in *ep300a* morphants but not in *crebbpa* morphants upon treatment with HDACi II, HDACIII and CHIC35. Lateral views with anterior to the top. Scale bar = 100 µm. Numbers at the bottom left corner represent number of embryos showing particular phenotype over the total number of embryos imaged.

### 2.6 RSTS patient-derived neural crest cells have a migration defect

The iPSC-derived neural model of Rubinstein Taybi Syndrome has been established to decipher the underlying mechanism of intellectual disability and for screening drugs to rescue the hyperexcitability that is observed in patient-derived neurons (24,25). Yet no RSTS iPSC-derived neural crest model has been generated to investigate the potential impact of *CREBBP/EP300* mutation on neural crest of RSTS patients. We generated an RSTS patient-derived iPSC line IGIBi018-A to test the effects on NCC migration (manuscript submitted).

We differentiated the RSTS patient-derived IGIBi018-A iPSCs into neural crest like cells to assess their migration potential in vitro using previously established protocols to differentiate iPSC into NCCs (26). Control iPSC (IGIBi19-A) and IGIBi018-A were differentiated into NCCs and compared for their migration potential and expression of neural crest markers p75 NGF and SOX10 (Fig.6 I-P). Neuroectodermal spheres were placed in fresh fibronectin-coated plates and time-lapse imaging was performed to observe the migration of the NCCs. By 5 hours post-adherence, control NCCs were seen migrating on to the substratum and by 10 and 15 hours, displayed extensive cell spreading (Fig. 6A-D, I-L). IGIBi018-A derived NCCs however, showed substantially less migration even at 15 hours (Fig. 6G-H, O-P). This suggests that in human NCC-like cells CREBBP is essential for efficient migratory behaviour and that this might be true in human RSTS embryos as well.

**Figure 6:**
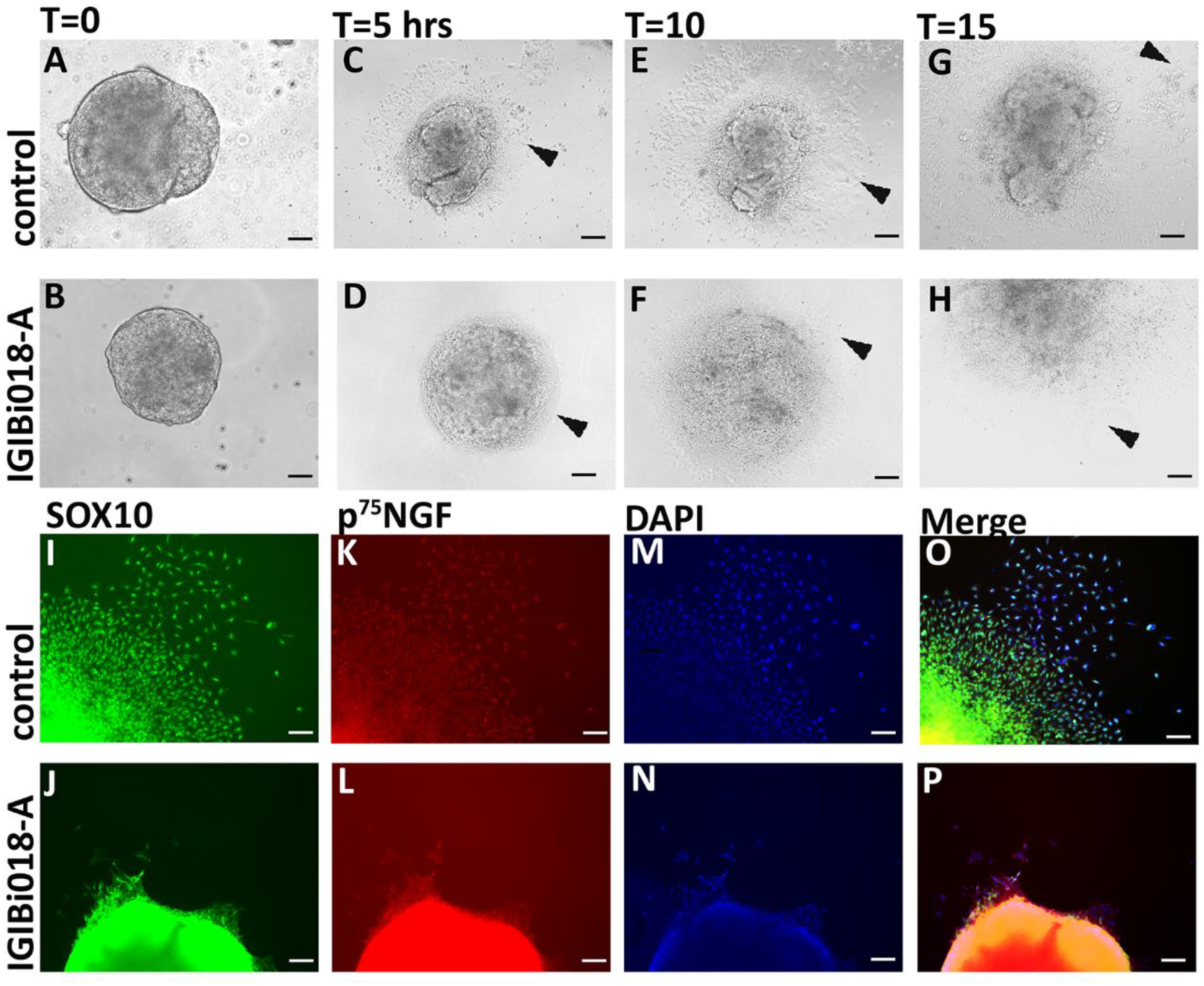
Migration defect in RSTS patient-derived Neural crest cells. **(A-H)** Brightfield images of NCC migration from the neuroectodermal sphere at various time points 0h, 5h, 10h and 15h **(I-P)** Immunofluorescence of neural crest markers SOX10 and p^75^NGF at T=15 hours in control and IGIBi018-A derived neural crest cells. Scale bar = 100 µm.

## 3. Discussion

Neural crest (NC) is a transient cell population that emerges from the epithelial neural tube during development to become mesenchymal and migratory. Neural crest cells harbor stem cell-like abilities and undergo several fate transitions throughout their life. These are cells that emerge from neuroectodermally committed origin to give rise to cells as diverse as cartilage, endocrine cells, melanocytes, neurons and glia (27,28). A number of gene regulatory networks are active during these fate transitions and epigenetic regulators play a very critical role in the emergence, migration and differentiation of neural crest (29).

The wide variety of tissues that trace their origin to neural crest also means that neural crest defects account for a large proportion of common birth defects (30,31). Diseases that result from defects in NC development are known as neurocristopathies; examples include Treacher-Collins syndrome (32,33), CHARGE Syndrome (26,34), Hirschsprung Disease (7), Bardet–Biedl syndrome (35). Rubinstein Taybi Syndrome has not been classified as a neurocristopathy yet [180849 (RSTS1), 613684 (RSTS2)] (1–3). However, many of the characteristic features of RSTS indicate that these could be linked to neural crest defects. RSTS patients have specific craniofacial indications such as broad nasal bridge, prominent beaked nose with deviated nasal septum, arched palate, and micrognathia (36); heart defects such as ventricular septal defect, atrial septal defect and patent ductus arteriosus (37) and severe constipation (5,14). A large proportion of RSTS patients have crowded and malpositioned teeth (38). RSTS patients reported with cases of neuroblastomas, tumors of neural crest origin (39–41).

With multiple indications that neural crest defects might play a central role in RSTS we asked whether the neural crest is affected in the zebrafish model of RSTS. We show that chemical inhibition of KAT3 protein activity with C646 in zebrafish embryos result in defects in craniofacial cartilage (11). In this study, we found that the knockdown of *ep300* and *crebbp* individually as well as together led to defects in craniofacial cartilage, melanophores, heart and enteric neurons, all tissues derived from the neural crest. Further, we discovered that C646-mediated inhibition of KAT3 proteins as well as knockdown of *ep300* and *crebbp* caused aberrant migration of neural crest cells from as early as 6 somite stage. This migration defect was also evident in 24 hpf embryos. Much stronger inhibition of neural crest migration was also found in CRISPR generated *ep300a* ^+/BD1Δ^ mutant fish embryos.

We reasoned that if a reduction of histone acetyl transferase activity was leading to neural crest migration defects in zebrafish embryos, inhibition of histone deacetylase activity may rescue the defects. We screened and found three such compounds, HDACi II, HDACi III and CHIC35, that could partially rescue the neural crest defects in the *ep300a* but not in *crebbpa* morphant embryos. This suggests that a global change in histone acetylation levels could specifically compensate for loss of ep300a HAT activity in neural crest cells, but not for crebbpa loss.

We generated an iPSC line (IGIBi018-A) from an RSTS patient with a mutation in the *CREBBP* gene to test if the neural crest defect is species specific (manuscript submitted). The IGIBi018-A cells were differentiated into the neural crest lineage in vitro. We observed that SOX10 positive neural crest cells in culture were defective in migration establishing that both EP300 and CREBBP have essential roles to play in neural crest migration in vertebrates.

So, how does KAT3 control neural crest migration? We identified gene enhancers occupied by EP300 in human neural crest cells in culture by analyzing publicly available data. These genes are putative effectors of the EP300 regulation in the neural crest. Our experimental analysis showed that the Epithelial to Mesenchymal transition (EMT) genes *snai1b* and *snai2* are downregulated in *ep300a* and *crebbpa* morphants. In the avian system *snai2* has been shown to repress *cadherin6b* (*cdh6b*) directly during EMT of neural crest cells (42). *Cadherin 6b* repression is essential for NC cells to delaminate and initiate migration (21). Downregulation of *snai2* in the *ep300a* and *crebbpa* morphants appears to cause de-repression of *cdh6b* in the zebrafish embryos. Thus, we propose that the KAT3 proteins Ep300a and Cbpa control neural crest migration by inducing the expression of EMT regulators *snai1b* and *snai2*. This in turn keeps *cdh6b* repressed facilitating migration of neural crest.

Our results suggest that many facets of Rubinstein Taybi Syndrome may be attributed to defective NC migration and here we make the case for classifiying RSTS as a neurocristopathy.

## 4. Material and methods

### 4.1 Zebrafish lines and Maintenance

Zebrafish (*Danio rerio*) were bred, raised and maintained at 28.5° C under standard conditions as described (43). Morphological features and timing (hours post fertilization, hpf) were used for staging the embryos as described previously (44). Embryos older than 24 hpf were kept in 0.003% 1-phenyl-2-thiourea (PTU) for RNA in-situ hybridization and fluorescence imaging. Zebrafish experiments were carried out according to the standard protocols approved by Institutional Animal Ethics Committee (IAEC) of CSIR-Institute of Genomics and Integrative Biology. Zebrafish lines used in this study are *Tubingen* (TU), *Tg(NBT:dsRed)*(45), *Tg(myl7:RFP*) (46) and *Tg(sox10:eGFP)* (47).

### 4.2 Chemical Treatment

Zebrafish embryos were treated with the compounds of interest in a 12-well plate at 11 hpf until the stage of interest. Embryos were then fixed with 4% paraformaldehyde and further processed for RNA insitu hybridization. Dimethyl sulphoxide (DMSO) was used as vehicle control since all the chemical compounds were dissolved in DMSO. The volume of DMSO given to control was equivalent to the maximum volume of the drug administered in that specific experiment. The compounds used in the study are, C646 (Sigma Cat.no: SML0002), HDACi II (Sigma Cat. no.: 382148), HDACi III (Sigma Cat. No.: 382149), and CHIC35 (Sigma Cat. No: C8742).

### 4.3 Whole mount RNA in situ hybridization

RNA in situ hybridization was performed as described previously (48). RNA in situ hybridization was done using probes for following genes, *sox10, crestin, tfap2a, snai2, snai1b, sox8, mycn, msxb, tfap2a, dlx2a* and *cadherin6*. 986bps fragment of *snai2* and 1132bps of *cadherin6* was PCR amplified from cDNA of 24hpf embryos and cloned in pCR®4-TOPO® vector. This cloned plasmid was used as template for in vitro transcription. Primer sequences are given in Supplementary material Table 3.

### 4.4 RNA sequencing analysis

We used publicly available ChIP-Seq data of EP300 used for identify active enhancer in hNCCs to delineate its potential targets. RNA seq data for ESC, NEC and NCC was analyzed to identify potential genes having a role in neural crest development (15). To identify potential EP300 target genes, enhancer coordinates bound by EP300 were used to assign genes located within 100kb of the transcription start site (TSS) using PAVIS (16) (Supplementary Data 1). Genes having two-fold increase in hNCC in comparison to hNEC were selected (Supplementary Data 2). Then 121 common genes which were upregulated in RNA Seq analysis and were potential targets of EP300 were selected (Supplementary Data 3).

### 4.5 Bright Field and Fluorescence Imaging

For bright field imaging of embryos Zeiss (Semi 2000C) bright field microscope was used. For fluorescent imaging Zeiss AxioScope A1 microscope was used. Images were captured by Zeiss proprietary software and were processed and analyzed using Adobe Photoshop.

### 4.6 Confocal imaging

For live imaging, embryos were first anaesthetized using 0.004% Tricaine for 1 min and then molded in molding media made using 0.75% low melting agarose (LMA)) (ThermoFisher scientific Cat. no.: 16520100) in Tricaine embryo water. Then embryo was placed on a slide followed by 0.75% LMA maintained at 40°C. The embryo was aligned using ZEISS light microscope until the agarose was completely polymerized. Further, the petri dish was filled with embryo water with tricaine and closed to prevent drying up by laser heat.

Live embryo imaging was performed using the LEICA SP8 confocal scanning microscope in 10X objective with the chamber temperature maintained at 28°C. For live imaging, xyz format was chosen followed by defining the duration and z-axis. Z-positions above and below the plane of focus were taken and an approximate z-size of about 200 µm was maintained for all embryos. Maximum projections covering all the Z-frames were done and the respective video/image is exported in mp4/TIFF format at 1024 X 1024-pixel size. For GFP embryos, Argon laser was used and HYD1 detectors were preferred with the gain adjusted to 100%.

### 4.7 Morpholino injection

KAT3 expression was knockdown in zebrafish embryos by injecting morpholino antisense oligonucleotides in one cell stage embryos. Two previously reported splice block morpholinos designed against *ep300a* were used (11). Morpholinos were injected at minimum effective concentration i.e., 4ng of *ep300a* MO1 and 8 ng of ep300a in one cell stage embryos. Two splice block morpholinos were used to knockdown *crebbpa* and *crebbpb*. The design of morpholinos is depicted in Supplementary Fig. S1. 4ng of *crebbpa* MO1 and MO2 and 8ng of *crebbpb* MO1 and MO2 were injected in one cell stage zebrafish embryos. A scrambled morpholino was used as a control in all the experiments. To equalize the concentrations of *crebbpa* and *ep300a* morpholinos with the double morpholinos, we introduced a control to both *ep300a* and *crebbpa* morpholinos. Sequence of MOs are provided in Supplementary material Table 1.

### 4.8 Knockdown PCR

For the knockdown PCR experiments, RNA was extracted from pool of 30 embryos at 1 dpf using trizol reagent (Takara), chloroform (Sigma Cat. No.: C2432), and isopropanol (Sigma Cat. No.: I9516). It was further followed by DNase treatment to remove DNA contamination. From this RNA, cDNA was prepared using an Invitrogen superscript IV kit. Knockdown of *ep300a, crebbpa* and *crebbpb* was confirmed by the detection of mis spliced RNA in the MO injected embryos. Primer sequences are given in Supplementary material Table 2.

### 4.9 CRISPR-based mutagenesis

sgRNA targeting the zebrafish *ep300a* gene was selected using the CHOPCHOP webtool using default parameters (49). GeneArtTM Strings™ DNA Fragments for bromo domain was used for in vitro transcription using T7 MEGAshortscript Transcription Kit (ThermoFisher Scientific Cat. no.: AM1334) according to manufacturer’s protocols. 100pg of sgRNA was injected along with 500 pg of spCAS9 protein (Takara). The F0 embryos were grown to adulthood and were then outcrossed with wild type to give rise to F1 animals. In the F1 generation, genomic DNA was isolated from fin clips of putative mutants and the target region was amplified using primers provided (Primer sequence provided in Supplementary material Table 4). PCR product was given for Sanger sequencing to confirm the mutant.

F1 mutant was crossed with wild type to give F2 generation and genomic DNA was isolated by fin clip method. The target region was PCR amplified and cloned in pCR®4TOPO® vector. Clones were subjected to Sanger sequencing to confirm mutations.

### 4.10 Neural crest differentiation

To differentiate RSTS derived iPSC into neural crest cells (NCCs), iPSC clusters were extracted followed by culturing in neural induction medium with growth factors including B27 Supplement (Gibco: 17504-044), N2 supplement (Gibco: 17502-048), bFGF (PeproTech Cat. No.: 100-18B), Insulin (Sigma Cat. No.: I3536) and EGF (PeproTech Cat. No.: AF-100-15) (26). Neuroectodermal spheres were formed by day 7-9 of differentiation. These neuroectodermal spheres were transferred to fibronectin coated plates to allow migration of NCCs. RSTS iPSC derived NC cells were tested for expression of NC markers.

### 4.11 Immunofluorescence (IF) staining

The cells were harvested and fixed with 4% Paraformaldehyde (PFA) for 15 minutes at room temperature. Subsequently, the cells underwent permeabilization with 1% Triton X-100 for 10 minutes at room temperature, followed by a 30-minute incubation in blocking buffer (composed of 5% bovine serum albumin with 0.1% Triton X-100 in PBS) at room temperature. The appropriate primary antibody diluted in PBS with 1% BSA, was then applied to the cells and allowed to incubate overnight at 4°C. Cells were then treated with the respective secondary antibodies for 1 hour at room temperature. For nuclei staining, DAPI was added and incubated for 15 minutes at room temperature. Subsequently, the stained cells were visualized, and images were captured using fluorescence microscopy (FLoid™ Cell Imaging Station and EVOS M5000). Details regarding the antibodies can be found in Supplementary material table 5.

## 5. Star Methods

| Key Resource Table |  |  |
| --- | --- | --- |
| Reagent/ Resource | Source | Identifier |
| **Antibodies** |  |  |
| Mouse Anti SOX10 | Invitrogen | Cat# 14-5923-82;<br>RRID:AB_2572892 |
| Rabbit p75 NGF Receptor Antibody | Thermo Fisher Scientific | Cat# BS-0161R;<br>RRID:AB_10856008 |
| Alexa Fluor™ 594 goat anti rabbit | Invitrogen | Cat# A11012;<br>RRID:AB_2534079 |
| Alexa Fluor™ 488 goat anti-mouse IgM | Invitrogen | Cat# A11001;<br>RRID:AB_2534069 |

| Chemicals |  |  |
| --- | --- | --- |
| C646 | Sigma | Cat# SML0002 |
| HDACi II | Sigma | Cat# 382148 |
| HDACi III | Sigma | Cat# 382149 |
| CHIC35 | Sigma | Cat# : C8742 |

## Authors’ contributions

SV designed, performed and analyzed the experiments. SD assisted in cloning and identifying mutations in the iPSC line. SG performed the zebrafish experiments. KC performed the confocal imaging. DS provided the control iPSC line. BP and SB provided resources. SPR oversaw reprogramming and differentiation of iPSC. CS conceptualized, designed, procured funding and coordinated the study. SV and CS wrote the manuscript, SD reviewed and edited the draft. All authors gave final approval for publication.

## Acknowledgements

We thank Anand Kishore Mukherjee and Mercy Rophina for the help in ChIP seq analysis. We thank Dr. Vivek T Natarajan, Dr. Beena Pillai and Dr. Shantanu Chowdhury for the valuable suggestions. We also thank Shabnam Sehrish and Menlum B. Lepcha for final proof reading.

