## Supplementary Material for "KAT3 mutations impair neural crest migration through EMT regulators *snai1b a*nd *snai2* in Rubinstein Taybi Syndrome"

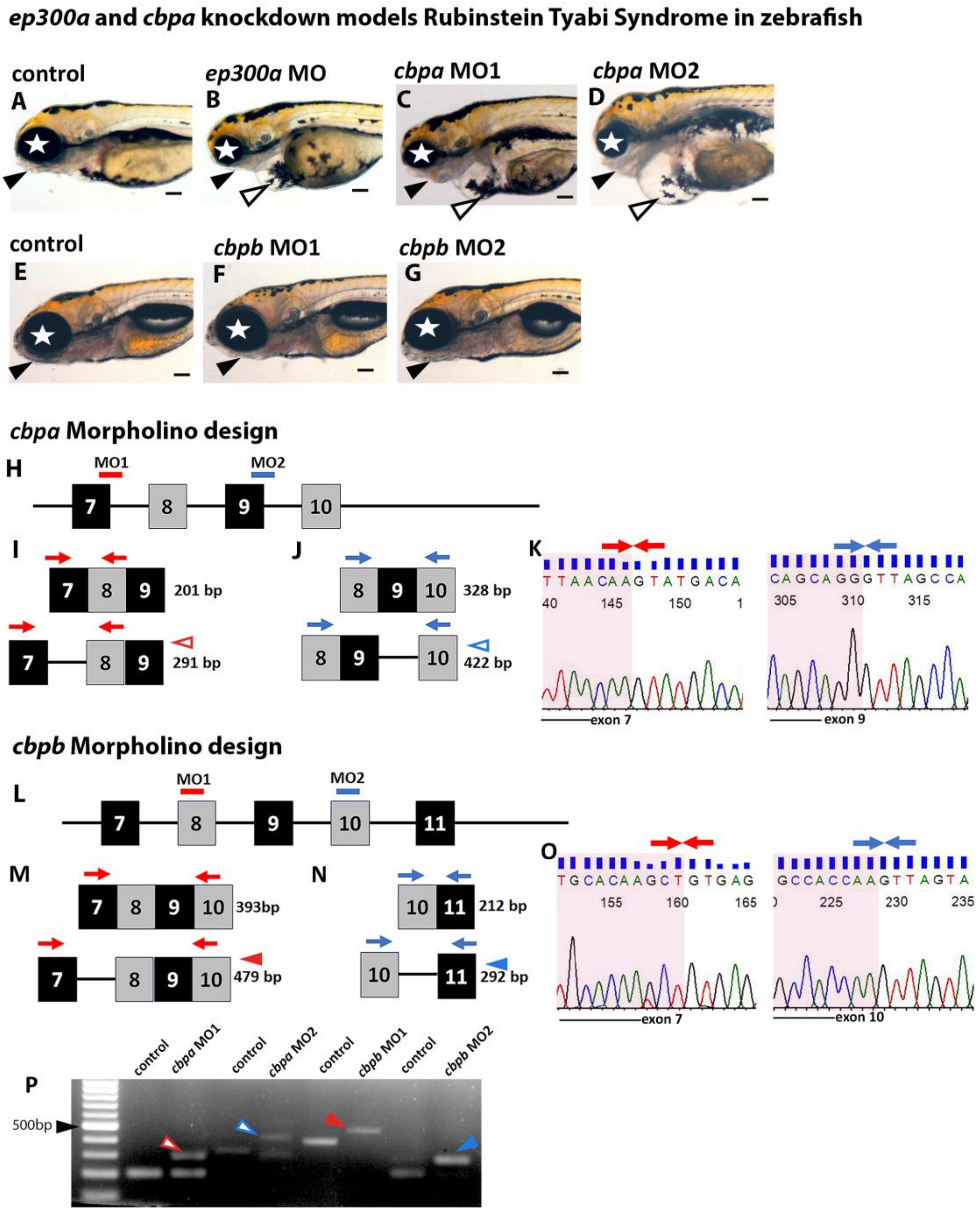


**Fig. S1. Knockdown of *crebbpa* and *crebbpb* using antisense morpholino morpholino oligonucleotides**

(A-D) 4dpf control larvae show well developed jaw (black arrowhead), normal heart (open arrowhead) and eye development (white star). Whereas, in *ep300a* and *crebbpa* morphants mandible is absent (black arrowhead), eye size is smaller (white star) and there is heart edema(open arrowhead).(E-G) *crebbpb* morphants show normal development and don't exhibit any defects. (H-O) Schematic representing the exon-intron splice site targeted by *crebbpa* and *crebbpb* morpholinos. Red and blue lines represent *crebbpa* MO1 and *crebbpa* MO2 respectively (Fig. S1 H, L). Intron retention due to splice block antisense morpholino oligonucleotides for *crebbpa* and *crebbpb* is shown in Fig.S1 (I, J, M, N). PCR was done to detect the retained intron due to mis-splicing. Expected PCR product size with intron retention is shown in (I) for *crebbpa* MO1(marked by red outlined arrow head), (J) for *crebbpa* MO2 (marked by blue outlined arrow head), (M) for *crebbpb* MO1(marked by red arrow head), (N) for *crebbpb* MO2 (marked by blue arrow head). (P) Agarose gel electrophoresis for amplicon from control and morpholino injected 24hpf zebrafish larvae. Intron retained PCR product corroborated with expected size (I, J, M, N). Amplicon with intron retention was sequenced. (K) Chromatogram show junction between exon7 and intron8 (red arrows) of crebbpa MO1 and junction between exon9 and intron 10 (blue arrows) of crebbpa MO2. (O) Chromatogram show junction between exon7 and intron8 (red arrows) of crebbpb MO1 and junction between exon10 and intron 11 (blue arrows) of crebbpbMO2.

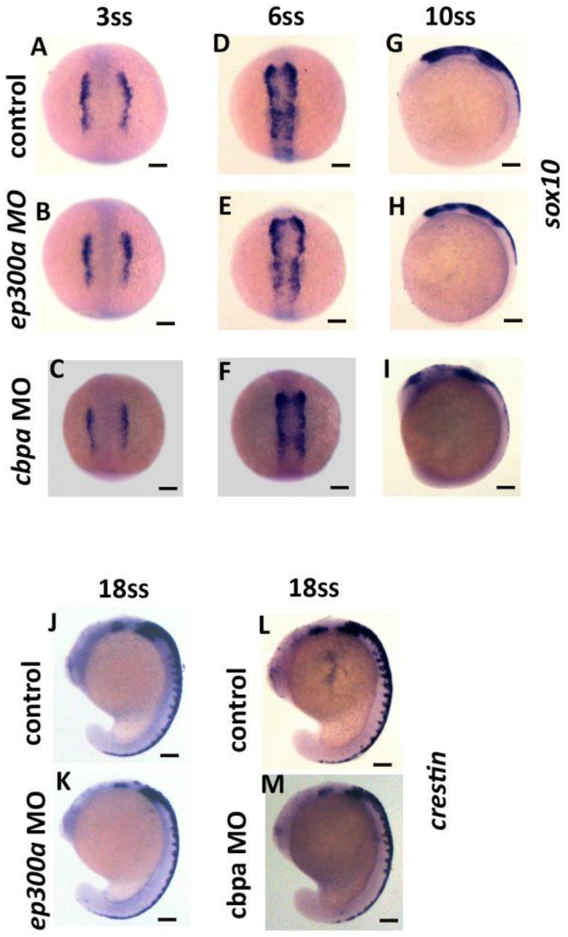


**Fig. S2. Effect of *ep300a* and *crebbpa* knockdown on neural crest development**

(A-I) RNA in-situ hybridization of *sox10* at various development stages of zebrafish. It shows that *sox10* expression remains unaltered in ep300a and crebbpa MO2 at (A-C) 3ss (D-F) 6ss (G-I) 10ss. (J-M) Expression of crestin at 18ss embryos stage. (J, L) control embryos show dorsoventral migrating neural crest cells in control (K, M) whereas these migrating neural crest cells are reduced in ep300a and crebbpa morphants. All images depict dorsal views with anterior to the left, and they are labelled with the time post-fertilization (hpf) or developmental stage. A scale bar of 100 μm is provided. The numbers in the bottom left corner denote the specific number of embryos out of the total represented in the image.

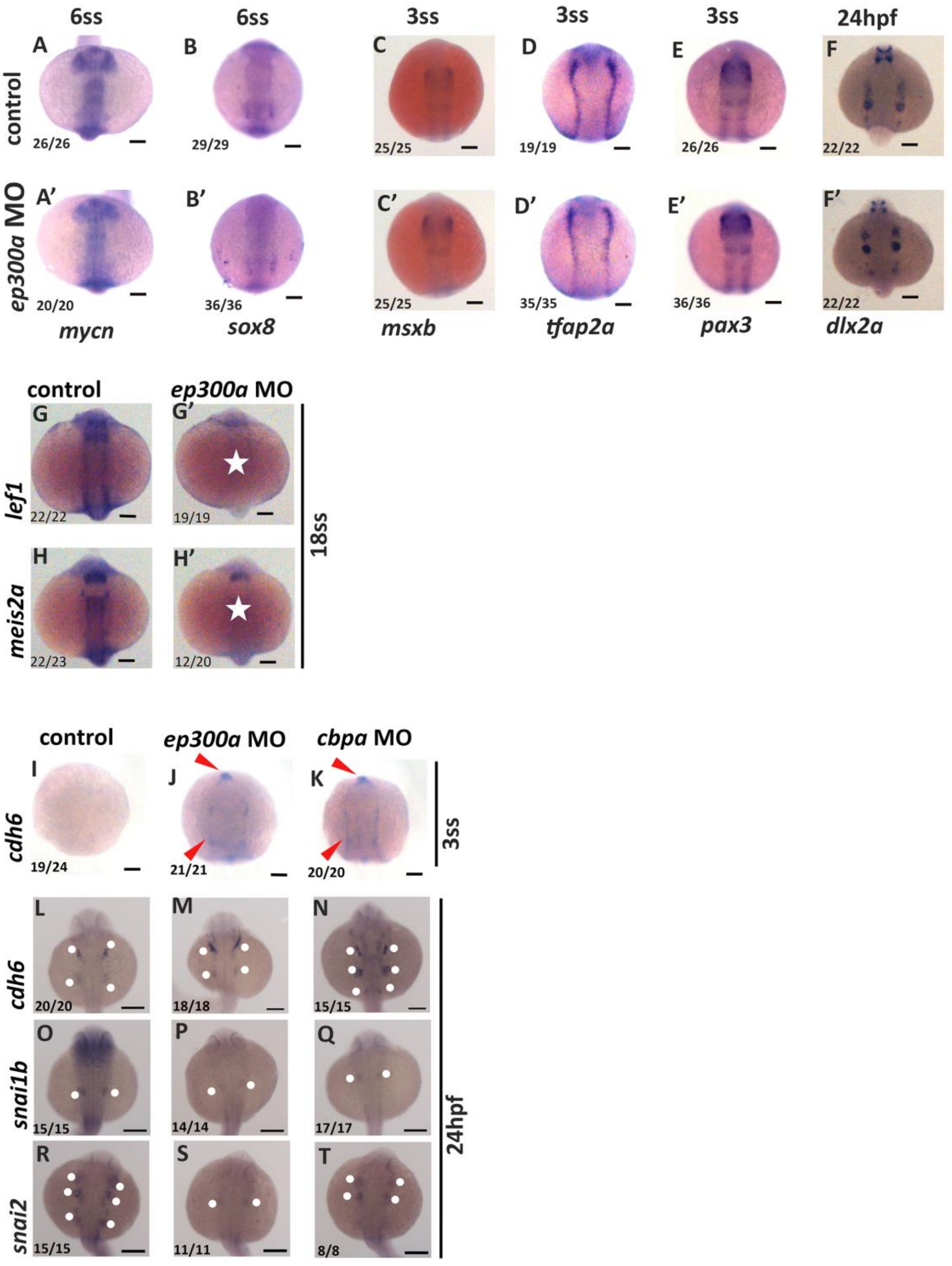


**Fig. S3 Effect of *ep300a* and *crebbpa* knockdown on putative effector genes involved in neural crest development**

(A-F') shows the expression of putative effector genes, *mycn*, *sox8*, *msxb, tfap2a, pax3* and *dlx2a* remain unchanged in *ep300a* morphants (G-H') shows that expression of *lef1* and *meis2a* is reduced in ep300a morphants (white star). (C-K) shows increased expression of cadhein6 in ep300a and crebbpa morphants at 3ss (red arrow heads). (L-T) *ep300a* and *crebbpa* morphants show increased expression of *cadherin6* and decreased expression of *snai2* and *snai1b* at 24 hpf (white dots). All images depict dorsal views with anterior to the left, and they are labelled with the time post-fertilization (hpf) or the developmental stage. A scale bar of 100 μm is provided. The numbers in the bottom left corner denote the specific number of embryos out of the total represented in the image.

**Supplementary Table**

Table 1: Morpholino sequence used in paper

| **Target gene** | **Sequence** |
|---|---|
| *ep300a* MO1 | 5'-GGGTCTGAACTATTACAAACCATGC-3' |
| *crebbpa* MO1 | 5'-GGCTTGTGTTTTCATGTCATACTTG-3' |
| *crebbpa* MO2 | 5'-ATTATCTTTCTGGCTAACCCTGCTG-3' |
| *crebbpb* MO1 | 5'-ATGTTTGATCTGTTGAACTCACAGC-3' |
| *crebbpb* MO2 | 5'-AAATACTGAACGGCTACTAACTTGG-3' |

Table 2: Primer sequence for Knockdown pCR

| **Gene** | **Primer** | **Seqence** |
|---|---|---|
| ep300a MO1 | Forward Primer | 5'-GACAAGAAGCCTCTGCCCAT-3' |
| ep300a MO1 | Reverse Primer | 5'-CACTGCCGTACTTCCCCATT-3' |
| crebbpa MO1 | Forward Primer | 5'-GGAGGTCAGTTGAACCCAATC-3' |
| crebbpa MO1 | Reverse Primer | 5'-GAGGTCCTGAGTGACGTGCT-3' |
| crebbpa MO2 | Forward Primer | 5'-ATGCCAGACGGCTCTACAGT-3' |
| crebbpa MO2 | Reverse Primer | 5'-GCAACCGTGACCTTCTCTTC-3' |
| crebbpb MO1 | Forward Primer | 5'-CGATCAGACGAACCTTCACA-3' |
| crebbpb MO1 | Reverse Primer | 5'-TTCGACAGCCTGGACTTTCT-3' |
| crebbpb MO2 | Forward Primer | 5'-TGAGTATTATCACTTTCTGGCTGA-3' |
| crebbpb MO2 | Reverse Primer | 5'-GGGCCATCAACTGATTTGG-3' |

Table 3: Primer sequence for in-situ riboprobe fragment

| **Gene** | **Primer** | **Sequence** |
|---|---|---|
| *snai2* | Forward | 5'-AAGATGAGGCTCTGCTGGAA-3' |
| *snai2* | Reverse | 5'-TGAAATCGTCTTTTGCGATG-3' |
| *cadherin 6* | Forward | 5'-GCGGAAAAGATGAGGACTTG-3' |
| *cadherin 6* | Reverse | 5'-CATCCACATCCTCGACACTG-3' |
| *mycn* | Forward | 5'-ATCGTGACtaatacgactcactatagggagaGAAGAGGCGTTCCATCACCA-3' |
| *mycn* | Reverse | 5'-atgtagctAATTAACCCTCACTAAAGGgagaGCGCAGAGGTACATCCATGA-3' |
| *sox8* | Forward | 5'-ATCGTGACtaatacgactcactatagggagaCTGAAGGGCTACGACTGGTC-3' |
| *sox8* | Reverse | 5'-atgtagctAATTAACCCTCACTAAAGGgagaGGCTCCTGCTTCTGCTTCAT-3' |

Table 4: Primer sequence for ep300 mutation confirmation

| **Gene** | **Primer** | **Sequence** |
|---|---|---|

| | | |
|---|---|---|
| *ep300a* | Forward | 5'-AAACGAAAGCTTGACACAGGC-3' |
| *ep300a* | Reverse | 5'-AGAATCAGCCCCAAACTCTCA-3' |

Table 5: Antibodies used for immunocytochemistry

| | **Antibody** | **Dilution** | **Catalog no.** |
|---|---|---|---|
| Primary Antibody | Mouse Anti SOX10 | 1:200 | Invitrogen Cat# 14-5923-82 |
| | Rabbit p75 NGF Receptor Antibody | 1:200 | Thermo Fisher Scientific Cat# BS-0161R |
| Secondary antibodies | Alexa Flour$^{TM}$ 594 goat anti rabbit | 1:500 | Invitrogen Cat# A11012 |
| | Alexa FluorTM 488 goat anti-mouse IgM | 1:500 | Invitrogen Cat# A11001 |
